# Microsaccades as a marker not a cause for attention-related modulation

**DOI:** 10.1101/2021.09.11.459890

**Authors:** Gongchen Yu, James P. Herman, Leor N. Katz, Richard J. Krauzlis

**Author notes:** Correspondence (G.Y.), (R.J.K.).

## Abstract

Recent evidence suggests that microsaccades are causally linked to the attention-related modulation of neurons – specifically, that microsaccades towards the attended location are required for the subsequent changes in firing rate. These findings have raised questions about whether attention-related modulation is due to different states of attention as traditionally assumed or might instead be a secondary effect of microsaccades. Here, in two rhesus macaques, we tested the relationship between microsaccades and attention-related modulation in the superior colliculus, a brain structure crucial for allocating attention. We found that attention-related modulation emerged even in the absence of microsaccades, was already present prior to microsaccades towards the cued stimulus, and persisted through the suppression of activity that accompanied all microsaccades. Nonetheless, consistent with previous findings, we also found significant attention-related modulation when microsaccades were directed towards, rather than away from, the cued location. Thus, in contrast to the prevailing hypothesis, microsaccades are not necessary for attention-related modulation, at least not in the superior colliculus. They do, however, provide an additional marker for the state of attention, especially at times when attention is shifting from one location to another.

## Introduction

The allocation of visual spatial attention is associated with both the enhancement of neuronal activity at the attended location (Desimone & Duncan, 1995; Krauzlis, Lovejoy, & Zenon, 2013; Reynolds & Chelazzi, 2004) and the tendency to make microsaccades towards the covertly attended stimulus while fixating (Engbert & Kliegl, 2003; Hafed & Clark, 2002; Lowet et al., 2018). A recent study provided evidence that the generation of microsaccades could play a causal role in the attention-related modulation of neuronal activity (Lowet et al., 2018). In a spatial attention task in which subjects were rewarded for making a saccade to a cued stimulus after it changed color, cortical neurons displayed attention-related enhancement only following microsaccades directed towards the attended stimulus.

These findings provide interesting evidence about the links between covert attention, neuronal modulation and fixational eye movements. However, they also raise potentially serious questions about how to interpret neuronal data obtained during the fixation tasks commonly used to study visual attention. Given that microsaccades are unavoidably generated during fixation, could the well-known modulation of neurons during covert visual attention tasks be an artifact of microsaccade generation? Is attention-related neuronal modulation obligately tied to the oculomotor intention to orient toward the attended stimulus, as suggested by the premotor theory of attention (Rizzolatti, Riggio, Dascola, & Umilta, 1987), and this dependence has been overlooked because sometimes it is a microsaccade towards the attended stimulus?

Here, we tested the relationship between microsaccade generation and attention-related neuronal modulation in the primate superior colliculus (SC), one of the most important brain structures for the control of visual spatial attention. Neurons in the SC provided the first evidence for neural correlates of visual attention (Goldberg & Wurtz, 1972) and are modulated during both overt and covert attention tasks (Herman & Krauzlis, 2017; Ignashchenkova, Dicke, Haarmeier, & Thier, 2004). Most relevant to our questions, SC activity is causally related to behavioral performance in attention tasks: inactivation causes attention deficits specifically for the affected location and conversely, microstimulation can selectively facilitate attention performance (Cavanaugh & Wurtz, 2004; Herman, Katz, & Krauzlis, 2018; Lovejoy & Krauzlis, 2010; Muller, Philiastides, & Newsome, 2005). Thus, if microsaccades are a necessary part of allocating attention, their causal role should be evident in the activity of SC neurons.

## Results

To investigate the possible role of microsaccades in attention-related modulation in the SC, we collected neuronal activity and eye movement data in two monkeys (Macaca mulatta) during a covert spatial attention task. SC extracellular activity was recorded with linear electrode arrays and eye position was measured with an infrared eye-tracker.

In the covert spatial attention task (figure 1a), head-fixed monkeys were required to release a joystick in response to a change in color saturation for the stimulus patch at the cued location and to withhold their response if the change occurred at an opposing foil location. Monkeys were required to maintain central fixation throughout the entire trial such that any allocation of attention to the cued stimulus patch was only done covertly.

**Fig. 1.**
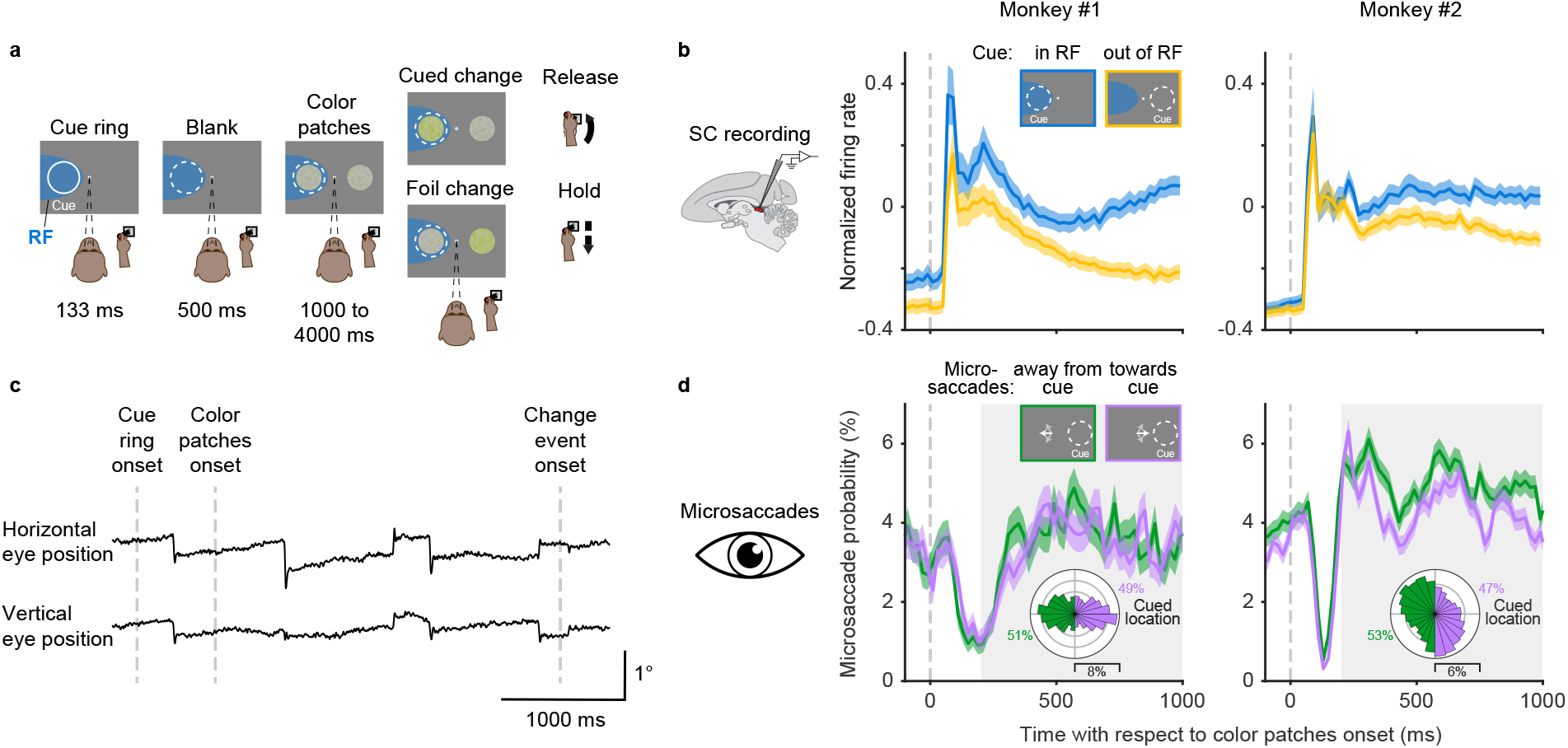
SC neuronal activity and microsaccades in a covert spatial attention task. **a**, The monkey was required to maintain central fixation, releasing a joystick in response to a color change at the cued location and holding their response if the change occurred at the opposing foil location. The dashed white ring illustrates the cued location and the blue shaded area denotes the response field (RF) of SC neurons; neither were visible to the monkey. **b**, Population SC average normalized firing rates for cue in RF (blue) and cue out of RF (yellow) conditions, aligned on the onset of the color patches. The insets illustrate the cue conditions when the SC RFs were on the left side. **c**, Single-trial horizontal and vertical eye position traces. The abrupt deflections are microsaccades. **d**, Session average probability of microsaccades toward the cued location (purple) and away from the cued location (green), aligned on color patches onset. The insets at top illustrate the microsaccade conditions when the cue was on the right side. The white arrows schematically represent microsaccades. The polar plots show the directional distribution of microsaccades during the delay period (grey shaded area), relative to the cued location. The numbers beside the polar plots denote the total proportion of microsaccades towards and away from the cued location. Error bars denote the standard error of the mean (SEM).

While monkeys performed the task, many SC neurons displayed attention-related modulation. The time course of SC population firing rate with respect to the onset of the color patches onset is shown in figure 1b for both monkeys (34 units in monkey 1 and 34 units in monkey 2). SC neurons displayed a phasic response shortly after the onset of the color patches, and during the delay period (200 to 1000ms) that followed, neuronal activity became steadily higher when the cued stimulus was inside the response field (RF) of the SC neurons compared to when it was outside the RF. This pattern recapitulates the well-known pattern of attention-related modulation in the SC and elsewhere (Desimone & Duncan, 1995; Herman et al., 2018; Herman & Krauzlis, 2017; Krauzlis et al., 2013; Reynolds & Chelazzi, 2004).

Simultaneous to SC neuronal recording, we measured the subjects’ fixational eye movements. Traces of horizontal and vertical eye position from one example trial are shown in figure 1c. Even though monkeys maintained fixation throughout the trial, they generated a series of fixational microsaccades consisting of abrupt and miniscule (typically smaller than 1 degree) deflections in eye position, as expected from previous findings (Hafed, 2011; Martinez-Conde, Otero-Millan, & Macknik, 2013; Rucci & Poletti, 2015).

To summarize when microsaccades occurred and where they were directed during the covert attention task, we calculated the probability of microsaccades on the same time axis as the SC neuronal firing rate (figure 1d). To identify how the direction of microsaccades was influenced by the location of the spatial cue, microsaccades were grouped based on whether they were directed towards or away from the cued hemifield. Overall, the probability of microsaccades both towards and away from the cued hemifield decreased sharply after patch onset and then rebounded during the following delay period. During the delay period (figure 1d, grey shaded area), the probability of microsaccades towards and away from the cued hemifield were similar, with slightly higher probability of away microsaccades in monkey 2. The polar plots in figure 1d provide a more detailed picture of the directional distributions of microsaccades relative to the cued location during the delay period. Microsaccades in monkey 1 showed two predominant directions directly towards and away from the cued location, whereas microsaccades in monkey 2 showed a weaker cue-related bias and idiosyncratic vertical tendencies unrelated to the cue. Thus, during sustained attention in the delay period, monkey subjects generated frequent microsaccades but their overall direction was not systematically biased towards the cued location.

### Effects of microsaccades on the time course of SC attention-related modulation

We next assessed how the occurrence of microsaccades was related to the time course of SC attention-related modulation (figure 2). Each panel in figure 2a depicts the population average normalized SC firing rate from subsets of trials in which there were either no microsaccades (top row), microsaccades towards the cued location (middle row), or microsaccades away from the cued location (bottom row), within a particular 100-ms epoch indicated by the gray shaded area. The eye speed trace in each panel serves to verify the inclusion or exclusion of microsaccades within each epoch. In total, we used 5 different epochs during the delay period, corresponding to the columns in figure 2a. To summarize the neuronal data, we measured the average normalized firing rates in 100-ms windows (dashed boxes) immediately after each microsaccade epoch and plotted these averages as a function of epoch in figure 2b. This choice of measurement window was motivated by the recent finding that cortical attention-related modulation emerged ~100ms after microsaccade onset (Lowet et al., 2018), but we found similar results with other windows (supplementary figure 1).

**Fig. 2.**
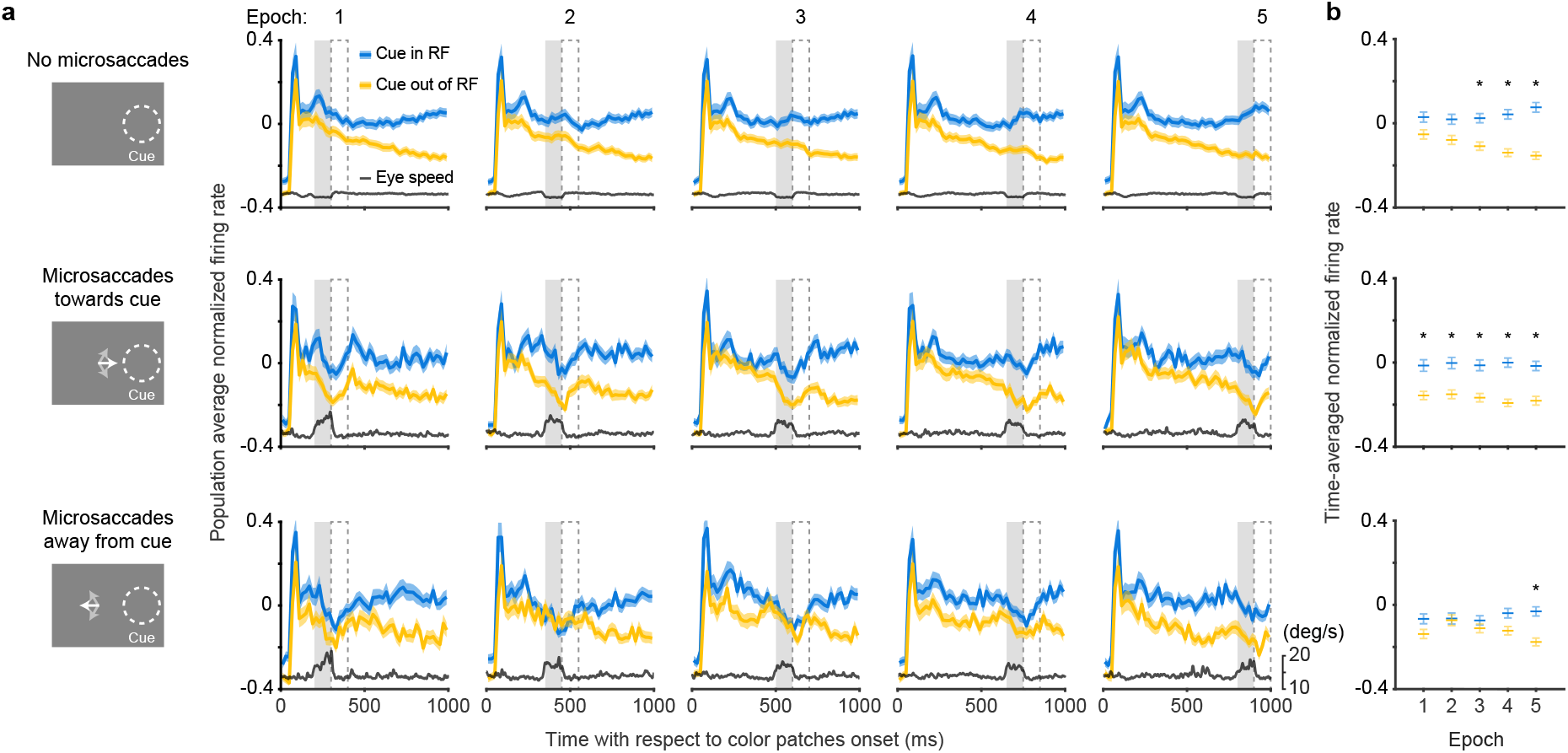
Effects of microsaccades on the time course of SC attention-related modulation. **a**, Each panel depicts the population SC average normalized firing rates (blue, cue in RF; yellow, cue out of RF) and eye speed traces (black) from subsets of trials in which there were either no microsaccades (top row), microsaccades towards the cue (middle row), or microsaccades away from the cue (bottom row), within a particular 100-ms epoch indicated by the gray shaded area. The chosen time epochs were: 200 to 300 ms, 350 to 450 ms, 500 to 600 ms, 650 to 750 ms, and 800 to 900 ms after color patches onset. The 100-ms dashed boxes following the gray shaded areas denote the time windows used for measuring time-averaged normalized firing rates for figure 2b. **b**, Time-averaged normalized firing rates as a function of epoch. The asterisks denote the epochs with significant higher activity for cue in RF than for cue out of RF, p < 0.05, ANOVA, Tukey-Kramer post-hoc comparisons. Error bars denote SEM.

In trials with no microsaccades in each epoch, the pattern of attention-related neuronal modulation (figure 2a, top row) was qualitatively similar to the one observed when including all trials (figure 1b). We found steadily higher population average neuronal activity during ‘cue in RF’ condition compared to ‘cue out of RF condition’ and this attention-related modulation was significant from the third epoch onwards (figure 2b, top row, ANOVA, post-hoc comparisons, p < 0.05). Thus, the exclusion of trials with microsaccades did not prevent the emergence of attention-related modulation over time during the delay period.

In trials with microsaccades in each epoch, SC neuronal activity was suppressed. Distinct dips in the firing rate traces were evident toward the end of each epoch, regardless of whether the microsaccades were directed towards the cue (figure 2a, middle row) or away from the cue (figure 2a, bottom row).

In addition to this suppression of firing rate that occurred regardless of microsaccade direction, we also found an attention-related modulation that did vary with the direction of the microsaccades. In trials with microsaccades towards the cued location (figure 2a, middle row), the difference between neuronal activity for ‘cue in RF’ versus ‘cue out of RF’ conditions was significant in all the epochs (figure 2b, middle row, ANOVA, post-hoc comparisons, all p < 0.05). In contrast, for trials with microsaccades away from the cued location (figure 2a, bottom row), the attention-related modulation was reduced and only emerged as significant in the last time epoch (figure 2b, bottom row, ANOVA, post-hoc comparison, p < 0.05).

Thus, the presence of SC attention-related modulation did not require the generation of microsaccades. Instead, microsaccades were associated with the suppression of firing rates, consistent with saccadic suppression (Hafed & Krauzlis, 2010). Nonetheless, there was a systematic relationship between the amplitude of the attention-related modulation and the direction of the microsaccades – we found more consistent attention-related modulation when microsaccades were directed towards the cue but not when they were directed away, suggesting that there might be a causal relationship between the two.

### Peri-microsaccade attention-related modulation

We next investigated the timing of the attention-related modulation relative to microsaccades by aligning SC neuronal activity to the onset of individual microsaccades directed towards (figure 3a, middle row) or away from the cued location (figure 3a, bottom row). For comparison, we also generated control data sets that contained no microsaccades during 400-ms windows chosen to match the timing of our microsaccade-aligned data (figure 3a, top row). In these ‘no microsaccades’ data, we still observed substantial attention-related modulation, again confirming that microsaccades were not necessary for SC attention-related modulation.

**Fig. 3.**
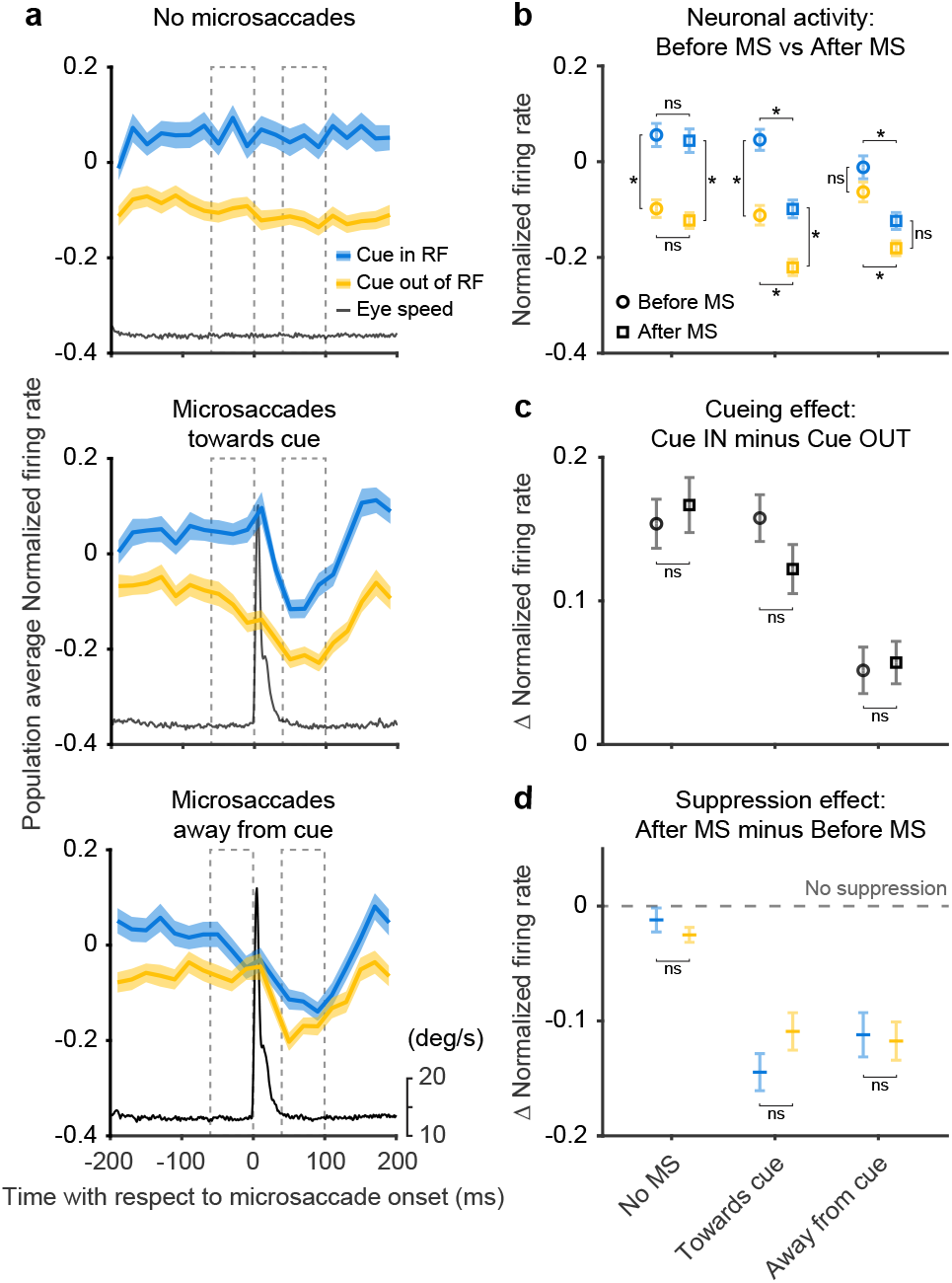
Peri-microsaccadic attention-related modulation. **a**, Population SC normalized firing rates aligned to the onset of individual microsaccades under 3 conditions: timing-matched no microsaccades (top row), microsaccades towards cue (middle row), and microsaccades away from cue (bottom row). The gray line denotes the average eye speed. The dashed boxes depict the windows we used to calculate the average normalized firing rates before (−60 to 0ms) and after (40 to 100ms) microsaccades. **b**, Average normalized firing rates before (circle) and after (square) microsaccades. **c**, The difference (Δ) in average normalized firing rates between cue-in-RF and cue-out-of-RF during ‘before microsaccade’ (circle) and ‘after microsaccade’ (square) windows. **d**, The difference (Δ) in average normalized firing rates between ‘after’ and ‘before’ window for cue in RF (blue) and cue out of RF (yellow) conditions. The dashed line indicates the level of ‘no suppression’. The asterisk denotes p < 0.05 and ‘ns’ denotes p > 0.05, ANOVA, Tukey-Kramer post-hoc comparisons. Error bars denote SEM.

For the data aligned on microsaccades, neuronal activity was suppressed immediately after microsaccade onset regardless of microsaccade direction or cue conditions (figure 3a), consistent with the results in figure 2. In contrast, the amplitude of the attention-related modulation did vary with microsaccade direction – the modulation was present only when the microsaccade was directed towards rather than away from the cued location. This result was robust with respect to the particular microsaccade inclusion window used for the analysis (supplementary figure 2).

The difference in attention-related modulation between microsaccade conditions (towards vs. away from cue) was not triggered by the microsaccade but was evident before microsaccade onset. To quantify changes in firing rates and attention-related modulation around the time of microsaccades, we calculated the average normalized firing rate before and after microsaccade onset (figure 3b). The dashed boxes in figure 3a depict the ‘before’ and ‘after’ windows we used for the calculation. For the ‘no microsaccades’ and ‘towards cue’ data sets, we found significant cue-related modulation (firing rate with cue in RF was higher than with cue out of RF) in both the ‘before’ and ‘after’ windows (ANOVA, post-hoc comparisons, all p < 0.05). In contrast, for the ‘away from cue’ data sets, we did not find significant attention-related modulation in either the ‘before’ or ‘after’ window (ANOVA, post-hoc comparisons, all p > 0.05). We also found a significant suppression of neuronal activity in the ‘after’ window compared to ‘before’, regardless of cue condition, for both microsaccade directions (ANOVA, post-hoc comparisons, all p < 0.05), but not for the no-microsaccade condition (p > 0.05).

As there were both attention-related modulation and response suppression around the occurrence of microsaccades, we investigated how these two processes interacted with each other. First, we tested whether the attention-related modulation present before microsaccades was changed when the overall neuronal response was suppressed after microsaccades (figure 3c). To compare the attention-related modulation, we measured the difference in population average normalized firing rates between cue in RF and cue out of RF conditions during both ‘before’ and ‘after’ windows. We did not observe any significant differences in attention-related modulation between ‘before’ and ‘after’ windows for any of the 3 microsaccade-related data sets (ANOVA, post-hoc comparisons, all p > 0.05). Thus, the amplitude of the attention-related modulation was largely preserved through the occurrence of the microsaccade.

Next, we investigated whether the microsaccade-related neuronal suppression observed in figure 3b was affected by the cueing condition (figure 3d). We calculated the difference in normalized firing rates between ‘after’ and ‘before’ windows for both ‘cue in RF’ and ‘cue out of RF’ conditions. We did not find any significant differences in the amplitude of the suppression effect across cueing conditions for any of the 3 microsaccade-related conditions (ANOVA, post-hoc comparisons, all p > 0.05). This indicates that the amplitude of the microsaccade-related suppression was not influenced by the cue-related modulation. Finally, we confirmed that our microsaccade-related findings could not be explained by motor effects related to the generation of microsaccades or variations in eye position (supplementary figure 3).

In summary, aligning firing rates on microsaccade onset clarifies the causal relationship between attention-related modulation and microsaccades. SC neurons display large attention-related modulation during epochs that contain no microsaccades. When microsaccades do occur, the attention-related modulation is present before microsaccade onset and has an amplitude at least as large as that found after microsaccades. Thus, microsaccades do not appear to cause the attention-related modulation but may be influenced by a shared process.

## Discussion

Our findings demonstrate that microsaccades are not necessary for attention-related modulation in the midbrain SC. First, attention-related modulation emerged over the same time course during the attention task, regardless of the generation of microsaccades (figure 2). Second, attention-related modulation was still observed when microsaccades were completely absent over long time periods (figure 3). Third, when microsaccades did occur, the neuronal attention-related modulation was already present prior to microsaccade onset and the occurrence of the microsaccade – and the accompanying saccade-related suppression – did not flip the pre-existing trend of attention-related modulation (figure 3). Thus, the attention-related modulation of SC neurons was readily dissociated from the occurrence of microsaccades during the sustained allocation of visual spatial attention.

Why are our findings different from those from a recent study showing that microsaccades play a causal role in the neuronal attention-related modulation (Lowet et al., 2018)? There are several factors that could have contributed to this difference – for example, differences in task design and brain regions. However, we think that the most salient factor is the time window used for microsaccade analysis. The previous study focused on microsaccades immediately after cue onset, which appeared after the visual stimulus was already present in the receptive fields of the neurons. Thus, the cue-induced change in attention state might be expected to trigger changes in both fixational eye movements and neuronal modulation. In our study, the cue was presented at the beginning of each trial and then disappeared (before the color stimuli were shown), and our analysis focused on a ‘delay period’ hundreds of milliseconds afterwards, which is the classic time window for studying sustained attention-related modulation. In future studies, it might be informative to directly compare the timing of attention-related modulation in different brain regions using these different cueing conditions.

Although our results demonstrate that microsaccades are not necessary for attention-related neuronal modulation, we did find that the amplitude of attention-related modulation varied with microsaccade direction. This dependence on direction preceded movement onset, consistent with previous behavioral work showing that microsaccades provide a marker for the state of attention (Engbert & Kliegl, 2003; Hafed & Clark, 2002; Yuval-Greenberg, Merriam, & Heeger, 2014). Finally, when using microsaccades as a marker to infer the state of attentive visual processing, one caveat raised by our study is that microsaccades are also obligatorily associated with suppression of neuronal activity, regardless of microsaccade direction and attention condition. Thus, before attributing changes in behavior or neuronal modulation following microsaccades to changes in the allocation of attention, the possibility that the changes are due to microsaccade-related suppression should also be considered.

## Materials and Methods

### General

We collected and analyzed data from two adult male rhesus monkeys (Macaca mulatta) weighing 9–12 kg. Data collection and analysis were not performed blind to the conditions of the experiments. All experimental protocols were approved by the National Eye Institute Animal Care and Use Committee, and all procedures were performed in accordance with the United States Public Health Service policy on the humane care and use of laboratory animals.

### Behavioral task

Monkey subjects performed a covert spatial attention task which has been described in detail previously (Herman et al., 2018; Herman & Krauzlis, 2017) and presented in figure 1a. In brief, monkeys initiated each trial by pressing down on a joystick, which triggered the presentation of a central fixation square. After monkeys acquired fixation, a white cue ring flashed (133ms) in the periphery indicating the ‘cued’ location. Then 500ms after the cue ring, two color patches were presented on the screen, with one patch in the same location as the previous cue ring (the ‘cued patch’) and another at equally eccentric location in the opposite visual hemifield (the ‘foil patch’). Within each trial, the mean saturation of one of the color patches could possibly change 1–4 s after color patches onset; if the change occurred at the ‘cued patch’, the monkeys were required to respond within 150 to 750ms by releasing the joystick, and if the change occurred at the ‘foil patch’ or if no change occurred, the monkeys needed to continue to hold the joystick down. In each block, cue-change, foil-change trials followed a ratio of 3:1 and all the trials were presented in pseudorandom order. The cue conditions were block designed (70 trials in each cued block) and the cued location alternated from block to block. The transition of cued block was indicated by single-patch trials (n = 18) with only one color patch inside the ‘cued’ location. These single-patch trials were also used for later control analyses (supplementary figure 3).

### SC recordings

We recorded SC extracellular activity in both monkeys using 24-channel or 32-channel V-Probes (50 μm spacing between contacts; Plexon Inc., Dallas, TX). In 3 sessions in monkey 1, we recorded both the left and right SC simultaneously. In 10 sessions in monkey 2, we used a single V-Probe to record in either the left or right SC. In each session, after advancing the V-Probe into the intermediate/deep layers of SC, we first monitored putative neuronal activity on each recording channel using a threshold-crossing method (μ ± 3σ on each channel). Based on the threshold crossing activity during a visually guided saccade task, we estimated the RFs for the neuronal activity on each recording channel. We then manually set the location of attention-task stimuli to be overlap with estimated RF centers. During the single V-Probe recording sessions in monkey 2, the two color patches were always 180° of elevation apart, with one patch inside the RFs. During the dual V-Probe recording sessions in monkey 1, the two color patches were placed at 0° and 200° of elevation to align stimulus location with both RFs.

Continuous electrophysiological data were acquired (40 kHz sample-rate) and high-pass filtered through an ‘Omniplex D’ system (Plexon Inc., Dallas, TX). Single units were sorted offline with Kilosort2 (Pachitariu, Steinmetz, Kadir, Carandini, & Harris, 2016). Because our intention was to test the relationship between SC attention-related modulation and microsaccade generation, we focused on SC neurons displaying classic visual attention-related modulation in our covert visual attention task which has been well established previously (Herman et al., 2018; Herman & Krauzlis, 2017). Neurons were identified as visually responsive if they displayed significant visual-evoked activity (50 to 150ms) after color patches onset compared to baseline (−100 to 0ms before color patches onset) in both single-patch and two-patch trials (p < 0.01, Wilcoxon rank sum test, two-sided). Visually responsive neurons with significant attention-related modulation during the later delay period (200 to 1000ms) were included for further analyses (mean firing rate for cue in RF was significantly higher than for cue out of RF, p < 0.01, Wilcoxon rank sum test, two-sided). In total, our dataset included 34 units in monkey 1 and 34 units in monkey 2.

### Microsaccade detection

Right eye position was monitored by an EyeLink 1000 infrared eye-tracking system (SR Research, Ottawa, Ontario, Canada) at a sample rate of 1000 Hz. Microsaccades were initially detected by using the 2D-velocity-based algorithm (relative velocity threshold = 4 and minimum saccade duration = 6 ms) developed by Engbert and Kliegl (Engbert & Kliegl, 2003). Every detected microsaccade was then inspected and visually verified by the experimenter. Microsaccade direction was calculated relative to the cued location (aligned in each session to 0 degree) in each trial. Microsaccades with directions ± 90 degree (window size 180 degree) relative to the cued location were grouped as ‘microsaccades towards the cued location’, and the other half of microsaccades were grouped as ‘microsaccades away from the cued location’.

### Firing rate analysis

Spike counts were binned in non-overlapping 20-ms windows. For normalization, each neuron’s spike counts was z-scored (mean subtracted and divided by standard deviation calculated from the neuron’s binned counts across trials and conditions).

### Timing match of microsaccades

When determining the attention-related modulation aligned to the onset of individual microsaccades (figure 3), we matched the timing of microsaccades and generated ‘no microsaccade’ control data. To do so, we first separated our analysis time period (200 to 1000 ms after color patches onset) into eight non-overlapping 100-ms bins, and computed the temporal distribution of trial counts for each microsaccade and attention condition. More specifically, for each bin we counted how many trials had microsaccades towards or away from the cued location, and how many trials had no microsaccades ± 200ms relative to the center of this bin. After counting all the trials, we used the lowest trial number within each temporal bin across conditions to subsample the data (without replacement) for all conditions, thereby matching the bin counts in the distributions across all conditions. For the ‘no microsaccade’ condition, the center of each bin was used to match the timing of microsaccades.

### Statistical analyses

To quantify how the occurrence or absence of microsaccades affected the time course of SC attention-related modulation (figure 2), we took the average normalized firing rates in 5 different time epochs during the delay period across all the conditions from each neuron (n = 68), and performed an ANOVA (total d.o.f. = 2039; error d.o.f. = 2010) with three factors: (1) time epoch (5 different time epochs, time epoch d.o.f. = 4), (2) attention conditions (cue in RF and cue out of RF, attention conditions d.o.f. = 1), and (3) microsaccade conditions (no microsaccades, microsaccades towards the cued location and microsaccades away from the cued location, microsaccade conditions d.o.f. = 2). We used Tukey-Kramer post-hoc comparison to test whether there was significant attention-related modulation (mean firing rate for cue in RF was significantly higher than with cue out of RF, p < 0.05) in each time epoch and each microsaccade-related condition.

To test the significance of attention-related modulation and firing rate suppression around the time of microsaccades (figure 3b), we computed the average normalized firing rate before and after microsaccade onset across conditions in each neuron (n = 68), and performed an ANOVA (total d.o.f. = 815; error d.o.f. = 804) with 3 factors: (1) time (before and after microsaccade onset, time d.o.f. = 1), (2) attention conditions (cue in RF and cue out of RF, attention conditions d.o.f = 1), and (3) microsaccade conditions (no microsaccades, microsaccades towards the cued location and microsaccades away from the cued location, microsaccade conditions d.o.f. = 2). The Tukey-Kramer post-hoc comparison was used to test whether there was significant attention-related modulation (mean firing rate for cue in RF was significantly higher than for cue out of RF, p < 0.05) and firing rates suppression (mean firing rate after microsaccade was significantly lower than before microsaccade, p < 0.05).

To test whether the attention-related modulation was significantly different before and after microsaccades (figure 3c), we calculated the difference in normalized firing rates between cue-in-RF and cue-out-of-RF conditions during both ‘before’ and ‘after’ windows in each neuron (n = 68), and performed an ANOVA (total d.o.f. = 407; error d.o.f. = 402) with 2 factors: (1) time (before and after microsaccade onset, time d.o.f. = 1) and (2) microsaccade conditions (no microsaccades, microsaccades towards the cued location and microsaccades away from the cued location, microsaccade conditions d.o.f. = 2). Tukey-Kramer post-hoc comparisons were used to test for significance (p < 0.05).

To test whether microsaccade-related neuronal suppression after microsaccades depended on cueing condition (figure 3d), we calculated the difference in normalized firing rates between the ‘after’ and ‘before’ windows for both cue-in-RF and cue-out-of-RF conditions in each neuron (n = 68), and performed an ANOVA (total d.o.f. = 407; error d.o.f. = 402) with 2 factors: (1) attention conditions (cue in RF and cue out of RF, attention conditions d.o.f. = 1) and (2) microsaccade conditions (no microsaccades, microsaccades towards the cued location and microsaccades away from the cued location, microsaccade conditions d.o.f. = 2). Tukey-Kramer post-hoc comparisons were again used to test for significance (p < 0.05).

To address whether peri-microsaccadic attention-related modulation was related to motor effects associated with microsaccade generation (supplementary figure 3), we performed control analyses using a subset of our data in which only a single patch was presented in the ipsilateral visual field, so that there was no visual stimulus inside the neurons’ RFs. We then aligned SC neuronal activity to the onset of microsaccades directed towards and away from the RF of the SC neurons. For comparison, we also analyzed firing rates from timing-matched epochs with no microsaccades. To test whether there were significant changes of activity around the onset of microsaccade, we took the average normalized firing in the ‘before’ and ‘after’ windows for each neuron (n = 68) and performed an ANOVA (total d.o.f. = 407; error d.o.f. = 402) with 2 factors: (1) time (before and after microsaccade onset, time d.o.f. = 1) and (2) microsaccade conditions (no microsaccades, microsaccades towards the RF and microsaccades away from the RF, microsaccade conditions d.o.f. = 2). Tukey-Kramer post-hoc comparisons were used to test for significance (p < 0.05).

To exclude the possibility that the peri-microsaccade attention-related modulation was related to the differences in eye position before microsaccades, we matched the eye positions distributions before microsaccades towards and away from the cued location across cue conditions (supplementary figure 3) and re-quantified the peri-microsaccade attention-related modulation. To match the eye positions, we first calculated the 2-D distribution of average eye position in the ‘before’ window using spatial bins of 0.25° × 0.25° across all conditions. We then used a subsampling procedure, like that described above to match the 2-D distributions across conditions. We then reanalyzed the average neuronal activity aligned on individual microsaccades for each neuron (n = 68) and performed an ANOVA (total d.o.f. = 543; error d.o.f. = 536) with 3 factors: (1) time (before and after microsaccade onset, time d.o.f. = 1), (2) attention conditions (cue in RF and cue out of RF, attention conditions d.o.f. = 1), and (3) microsaccade conditions (microsaccades towards the cued location and microsaccades away from the cued location, microsaccade conditions d.o.f. = 2). Tukey-Kramer post-hoc comparisons were used to test whether there was significant attention-related modulation (p < 0.05).

## Acknowledgements

We thank Nick Nichols, Daniel Yochelson, Denise Parker and Amber Lopez for technical support. We thank Xuefei Yu, Lupeng Wang, Christian Quaia, Sheridan Goldstein, Kerry McAlonan and Kara Cover for helpful discussions. This work was supported by the National Eye Institute Intramural Research Program at the National Institutes of Health (ZIA EY000511).

## Author contributions

G.Y., J.P.H. and R.J.K. designed the experiments. G.Y., J.P.H. and L.N.K. conducted the experiments. G.Y. and J.P.H. analyzed the data. G.Y. and R.J.K. drafted manuscript. All authors edited and revised manuscript.

## Supplementary Figures

**Supplementary Fig. 1.**
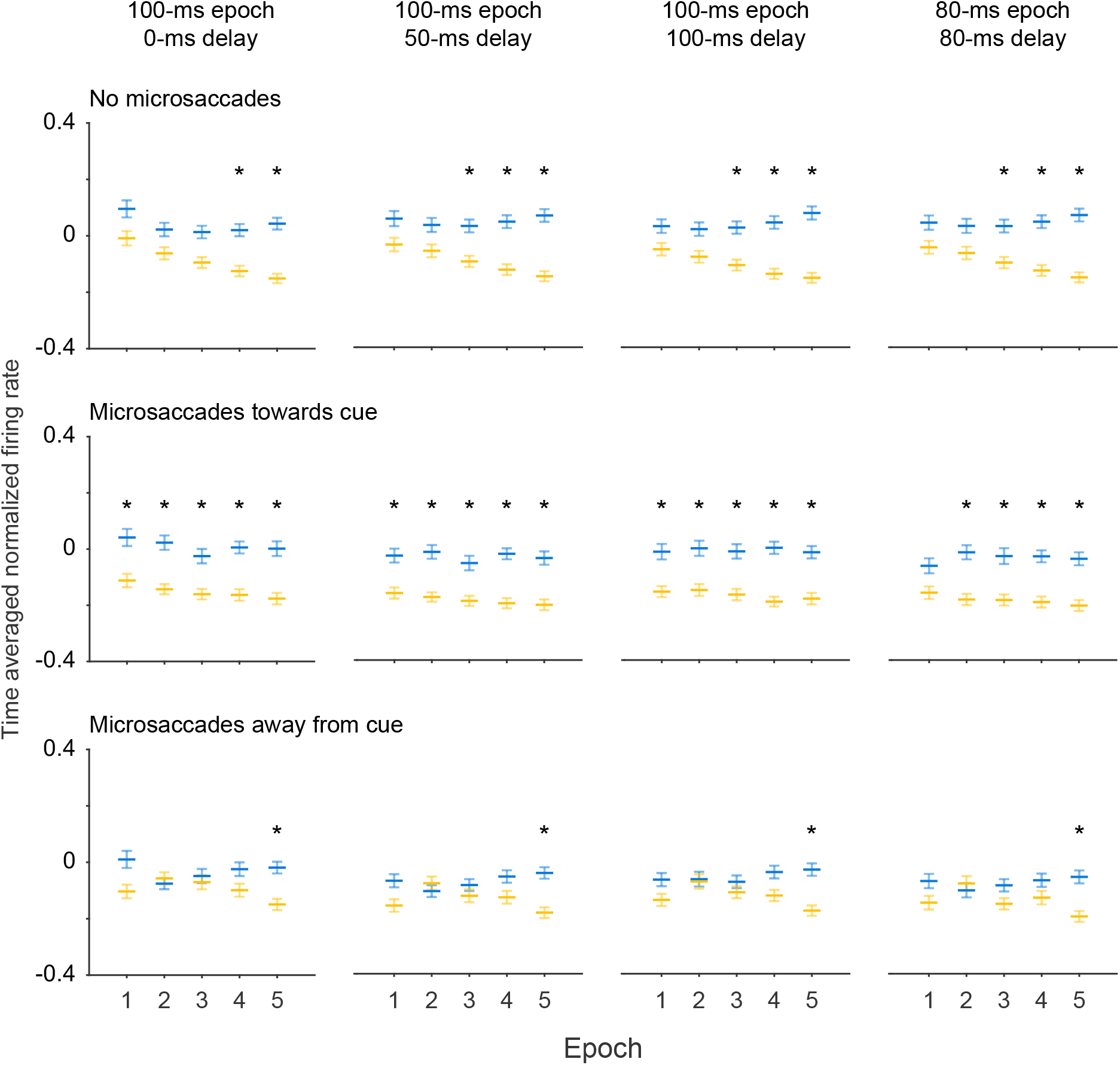
Effects of microsaccades on the time course of SC attention-related modulation were similar when using other measurement windows. In our main results, we used 5 different 100-ms epochs during the delay period to categorize trials based on the occurrence of microsaccades, and corresponding 100-ms windows (delayed by 100ms) for each epoch to measure the average neuronal activity. Here the plots are in similar format as figure 2b with no microsaccades (top row), microsaccades towards the cue (middle row) and microsaccades away from the cue (bottom row) conditions. But we tested 4 different combinations of microsaccade epochs and measurement windows: 100-ms microsaccade epochs with 100-ms measurement windows after a 0 ms delay, (first column), 100-ms epochs with 100-ms measurement windows after a 50 ms delay, (second column), 100-ms epochs with 100-ms measurement windows after a 100 ms delay, (third column, same as figure 2b), and 80-ms epochs with 80-ms measurement windows after an 80 ms delay (forth column). We observed qualitatively similar results across all combinations of delay and window duration. The asterisks denote the epochs with significant higher activity for cue in RF than for cue out of RF, p < 0.05, ANOVA, Tukey-Kramer post-hoc comparisons. Error bars denote SEM.

**Supplementary Fig. 2.**
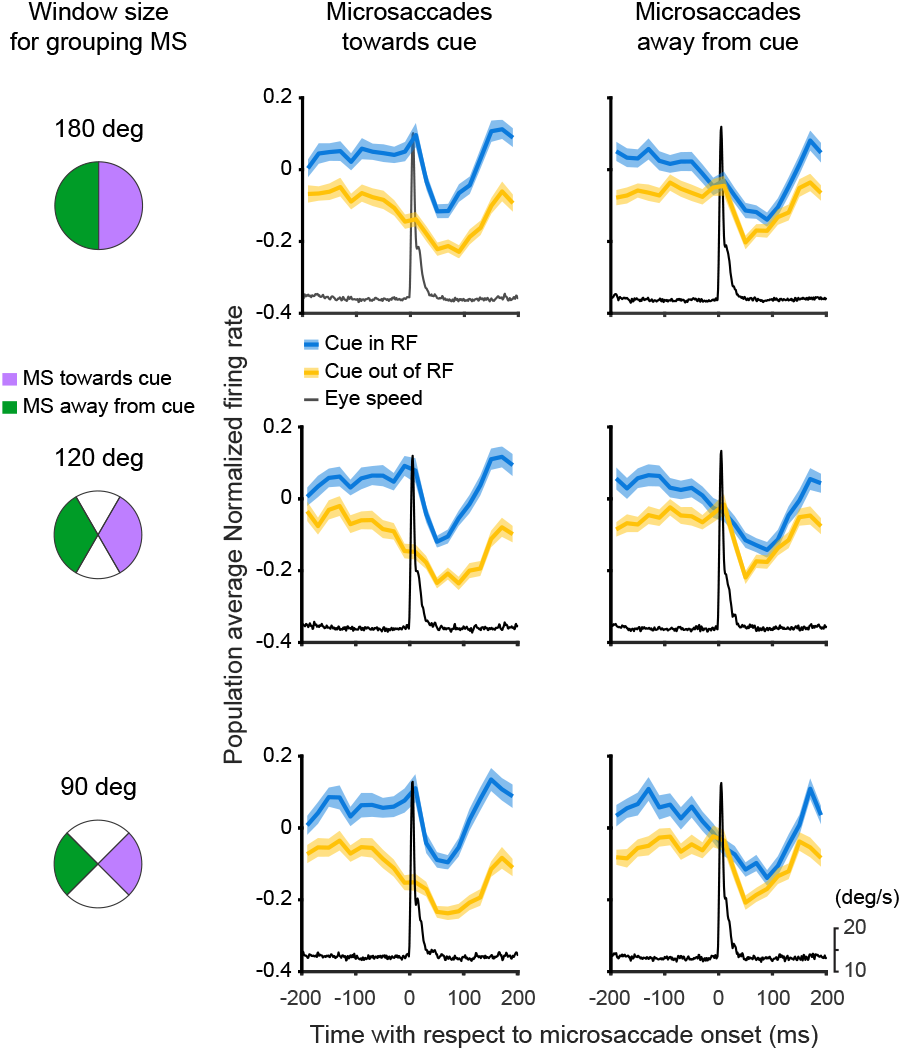
Peri-microsaccadic attention-related modulation was similar when using narrower windows to group microsaccades towards/away from the cued location. In our main results, microsaccades with directions ± 90 degree (window size 180 degree) relative to the cued location were grouped as ‘microsaccades towards the cued location’, and the other half of microsaccades were grouped as ‘microsaccades away from the cued location’. Here we tried three different window sizes to group microsaccades towards/away from the cued location: ± 90 degree (window size 180 degree, top row), ± 60 degree (window size 120 degree, middle row) and ± 45 degree (window size 90 degree, bottom row). We observed qualitatively similar results across the three window sizes.

**Supplementary Fig. 3.**
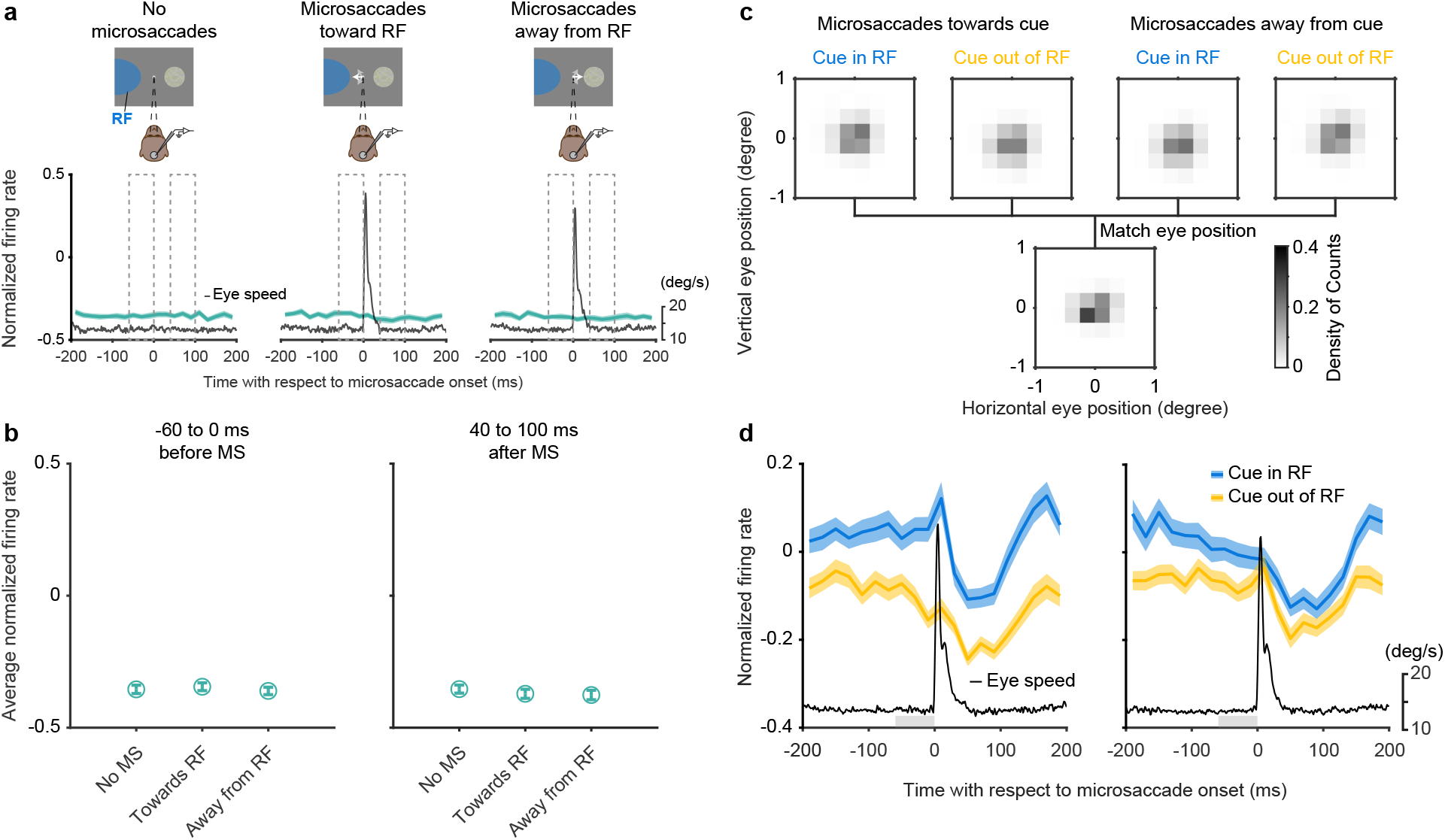
Peri-microsaccadic attention-related modulation was not explained by motor effects or differences in eye position. **a**, To address whether peri-microsaccadic attention-related modulation might be related to motor effects associated with microsaccade generation, we used a subset of our data in which only a single color patch was presented, and this single patch was located in the ipsilateral visual field (schemes above data panels), so that there was no visual stimulus inside the neurons’ RFs (blue shaded area) and any potential modulation would be due to motor effects, not sensory. The white arrows in the schemes denote microsaccades. SC normalized firing rates are plotted aligned on the onset of individual microsaccades (data panels) in each of three conditions: timing-matched no microsaccades (left), microsaccades towards SC RF (middle), and microsaccades away from SC RF (right). The gray line denotes the average eye speed. The dashed boxes depict the windows we used to calculate the time-averaged normalized firing rates ‘before’ (−60 to 0ms) and ‘after’ (40 to 100ms) microsaccades. **b**, Average normalized firing rates before microsaccades (left) and after microsaccades (right). Error bars denote SEM. If our SC neurons were modulated by the motor preparation of microsaccades, we should find higher activity for microsaccades directed towards the RF compared to those directed away, or the ‘no microsaccade’ dataset. Instead, we found no differences in activity across any of these conditions (ANOVA, post-hoc comparisons, all p > 0.05). Thus, the peri-microsaccade attention-related modulation cannot be explained by the motor effects of microsaccades. **c**, To address the possibility that the peri-microsaccadic attention-related modulation might be explained by systematic differences in eye position before microsaccades, we calculated the 2-D spatial distribution of eye positions (upper row) before the onset of microsaccades (calculation window indicated by the gray bars in panel b) across microsaccade directions (towards/away from the cue) and attention conditions (cue in RF/cue out of RF), matched all the distributions (lower row). The gray scale bar denotes the magnitude of the proportion. **d**, After matching eye position distributions, we recalculated the SC normalized firing rates (for cue in or out of the RF) aligned on the onset of individual microsaccades for conditions with microsaccades towards the cue (left) and microsaccades away from the cue (right). The gray trace indicates the eye speed. The light gray bars immediately above the horizontal axes depict the time windows (−60 to 0ms) used to match eye position distributions. The results were unchanged – the attention-related modulation was present only around microsaccades towards cued location (ANOVA, post-hoc comparison, p < 0.05) but not microsaccades away from cued location (ANOVA, post-hoc comparison, p > 0.05). Therefore, the peri-microsaccade attention-related modulation cannot be explained by differences in eye position before the microsaccade.

